# Modeling Human Peripheral Myelin Fully from Pluripotent Stem Cells and Immune-mediated Neuropathy *In Vitro* using ZIKA Virus

**DOI:** 10.1101/2020.05.23.112748

**Authors:** Fahimeh Mirakhori, Cheng-Feng Qin, Zhiheng Xu

## Abstract

The generation of *in vitro* model of human peripheral myelin development and associated disease from human pluripotent stem cells (hPSCs) has been a challenge so far. In addition, the underlying mechanism for ZIKA virus (ZIKV) infection incurred Guillain-Barré syndrome (GBS) remains unexplored due to the lack of a suitable model. Here, we report the *de novo* generation of a human peripheral myelination model with competent Schwann cells (SCs). Those human SCs generated from hPSCs via compound screening were capable of forming compact myelin both *in vitro* and *in vivo*. We found ZIKV infection caused GBS-like events *in vitro* including myelin sheath degeneration, as well as dysregulated transcriptional profile including the activated cell death pathways and cytokine production. These effects could be partially reversed by several pharmacological inhibitors. Our model therefore provides a new and robust tool for studying the pathogenic mechanisms and developing of therapeutic strategies for related neuropathies.

## Introduction

Peripheral neuropathies are myelination disorders and common neurological complications that can be caused by various conditions such as inflammatory/infectious diseases, diabetes, chemotherapy and aging or genetic mutations. These diseases show marked myelin sheath impairments, lead neurological deficits with severe clinical motor and/or sensory problems (Campana, 2007), affects 4-7% of the population (~20 million Americans) (Colloca et al., 2017). Currently, no effective treatments for those diseases are available mainly due to the lack of proper models, especially the humanized model for peripheral myelination (human neurons with human Schwann cells (SCs), the peripheral myelinating glial cells) to unravel the pathogenic mechanisms and to develop therapeutic strategies (Campana, 2007; Clark et al., 2017). Although brain organoid derived from pluripotent stem cells (hPSCs) has been widely applied to understanding stem cell biology, organogenesis, and various brain disorders (Amin. 2018), the generation of *in vitro* model of human peripheral myelin development and associated disease from hPSCs has not been successful. All available cellular models are based on interspecies models using rodent dorsal root ganglia (DRG) neurons or rodent SCs to co-culture with human pluripotent stem cell (hPSC) derived neurons or hPSC-Schwann cells (SCs). There is no analogous model available for human Myelination due to inability of hPSC-SCs to myelinate hPSC-neurons (Clark et al., 2017; Kim et al., 2017).

There are serious immune-mediated neurological complications associated with ZIKA virus (ZIKV) in adults such as Guillain-Barré syndrome (GBS, a post-infectious immune-mediated peripheral neuropathy) (Nascimento and da Silva, 2017). Although the peripheral neuropathy is well-documented, its underlying mechanism remains unclear due to the lack of an appropriate model. In this study, we endeavored to establish a humanized *in vitro* system of peripheral myelination consisting of a potent myelinating SC progenitor (SCp) population along with sensory neurons all generated from hPSCs. We developed a new hPSCs differentiation protocol for the generation of myelinated fibers via compound screening. We verified the peripheral myelination system with an expandable and functional SCs, capable of forming compact myelin both *in vitro* and *in vivo*. Moreover, we found that ZIKV infection of humanized peripheral myelination culture resulted in the activation of cell death pathways, alternation of transcriptional profile, cytokine over production and myelin sheath degeneration. These effects could be partially reversed by several pharmacological inhibitors. Our *in vitro* human myelination model with competent SCs provides a new and robust approach for studying the pathogenic mechanisms and developing of therapeutic strategies for related neuropathies.

## Methods

### Stem Cell culture and PNS differentiation

Human embryonic stem cells (hESC), line H9 (WA09, WiCell, USA-Passage 35-50), CRISPR/Cas9 modified H9 cell lines for AAV::EGFP, SOX10::EGFP and OCT4::EGFP cell lines(Oh et al., 2016), and hiPSC lines(Oh et al., 2016), passage 35-50) were cultured on irradiated mouse embryonic fibroblasts (MEFs-GlobalStem) in DMEM/F12, 20% Knocked out serum replacement (Invitrogen), 0.1 mM MEM-NEAA, 2mM L-glutamine, and 55 uM 2ME (LifeTechnologies) as KSR media, supplemented with (10 ng/ml) human recombinant FGF2 (R&D). Cells were monitored every 2 weeks for Mycoplasma and confirmed as Mycoplasma negatives for downstream usage.

For PNS differentiation, cells were harvested by Accutase (Invitrogen) and plated on 0.1% gelatin coated plates for 30 min at 37°C to exclude MEF contamination. Non-adherent cells were plated as single cells onto Geltrex (Invitrogen) coated 24 well plates at density of (2.4x 10^6^ cells/plate) in MEF condition medium with (10ng/ml) FGF2 and (10uM) Rock inhibitor (Y-27632 - Cayman chemical Co). Culture were allowed to expand in this condition until 60-70% confluent, at which time neural induction was started according to a modified version of LSB3i protocol developed by Lee et al., (2007) and Chambers et al. (2012). Briefly, media was changed gradually into a gradient of (1/4) Neurobasal media (NB: based medium contained 1% Lglutamine, 1% N2 and 2% B27 (all from Invitrogen)) plus reduced gradient of (3/4) KSR media every other day: For the first two days, this media was supplemented with (0.5 uM) LDN-193189 (Abcam), and (10 uM) SB431542 (Cayman). Followed by switching to (1/2) NB, (1/2) KSR with LDN, SB, (10uM) DAPT (Cayman) and (3uM) CHIR (Cayman) media for another 2 days. Cultures were kept in (3/4) NB, (1/4) KSR media plus DAPT, CHIR small molecule inhibition for 2 days more. After this step media was changed to (100%) NB supplemented with (200 uM) Ascorbic acid (Sigma) and (200 uM) dbcAMP (Sigma) for additional 12 days. Then, media was changed to a NB plus (200 uM) Ascorbic acid (Sigma), (50 ng/ml) human NGF, (20 ng/ml) BDNF, and (10 ng/ml) GDNF (all from PeproTech), for the next 3 weeks (referred as M1). For myelination step, we did drug screening to find the optimal condition (see Supplementary information, Fig. 1 and Table 1). Small molecules were diluted in DMSO in a way that its overall concentration in medium was kept throughout the experiments below 0.1%. Briefly, cells were exposed to a M1 media contained (10 ng/ml) human HRG1 (R&D) and (1uM) Clemastine Fumarate D3:D20 (Cayman) for 6 weeks (M2; the optimal media) and (2.5ug/ml) Laminin (Sigma), and (1 ug/ml) Fibronectin (R&D). We referred these 12 week-culture as mixed culture.

**Figure 1.**
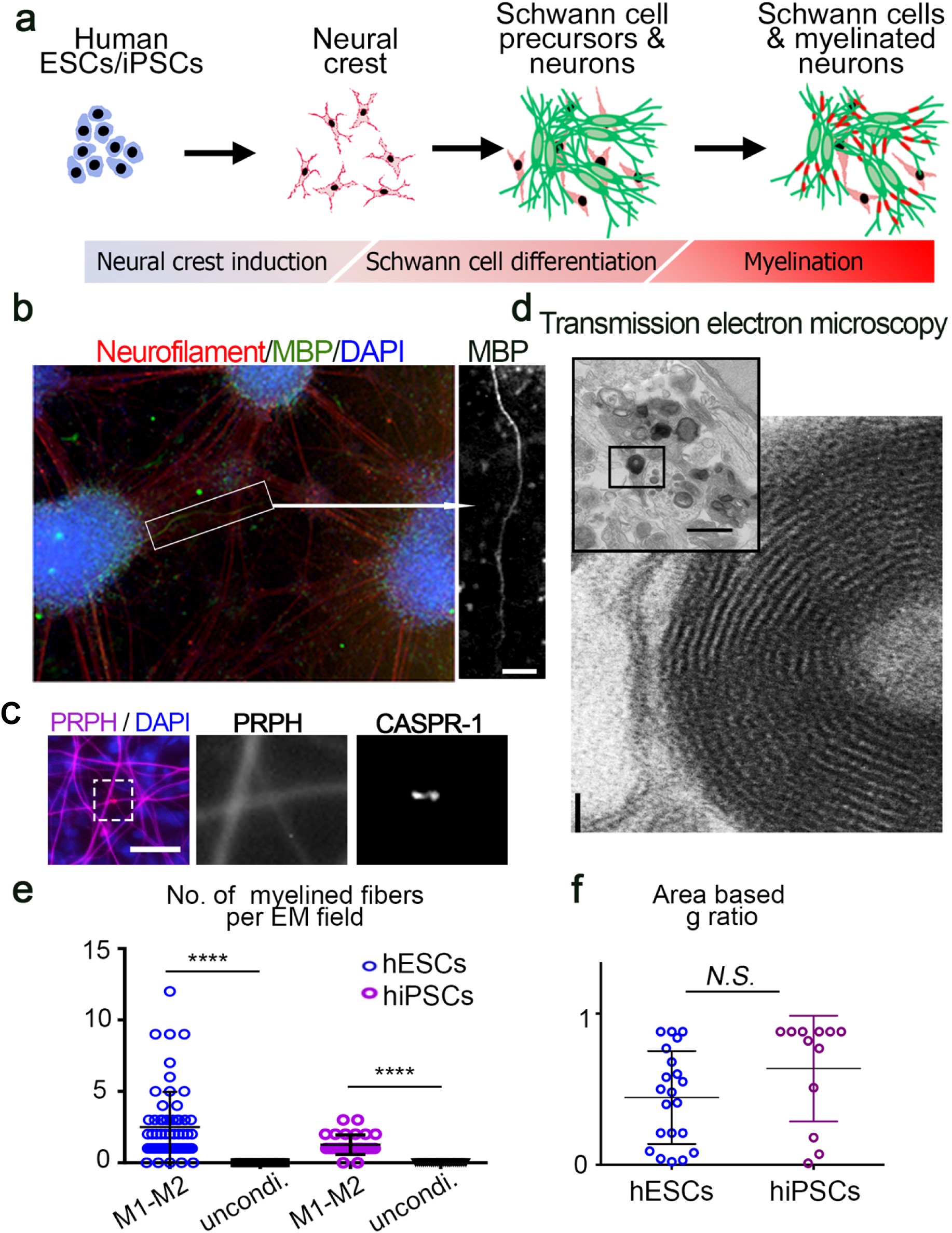
Development of an *in vitro* PNS myelination model from human PSCs. a. Schematic of 12 weeks peripheral neurons and Schwann cell differentiation protocol as mixed culture (for more details see also methods). b. Representative images for MBP and TUJ1 staining in the 12 weeks mixed culture (scale bars, 200 μm). c. Representative images for CASPR1 and Peripherin staining in the 12 weeks mixed culture (Scale bar, 50 μm in left panel), and the area of white dashed box in left panel is magnified in center and right panels. d. Ultrastructure of myelinated fibers. The myelination structure in boxed region of the upper left panel (scale bars, 100 μm) is shown in lower right panel (scale bars, 500 μm) e. Quantification of the number of myelinated fibers. M1, M2 condition refer to media 1 and 2 during differentiation, and unconditioned as control media; NB media plus DMSO, without additional factors. f. The area based g-ratio in human PSCs after 12 weeks *n* = 60 (f), *n* = 21 (f) for human ESC-derived cultures; *n* = 30 (e), *n =* 12 (f) for human iPSC-derived cultures. Data represent mean ± S.E.M., **** *P* < 0.0001, N.S. is non-significant, calculated by unpaired *t*-test (E, F).

### Immunofluorescence and antibodies

Cells were fixed with 4% paraformaldehyde for 30 min, washed with PBS, permeabilized using 0.1% Triton X100 in PBS and blocked using 1% BSA, 10% secondary host serum in PBS. The following primary antibodies were used: NF (DSHB and Sigma, 1:400), TUBB3 (Covance 1:400), MBP (Abcam, 1:50 and Covance, 1:100), CASPR-1 (a gift from Prof. Bhat, 1:400), MAG (Abcam and Sigma, 1:200), S100B (DAKO, 1:400), BRN3A (Millipore, mab1585, 1:500), GFAP (DAKO, 1:200), MPZ (Novus Biologicals, 1:200), NS1 (Oneworld lab, 1:200), Caspase3 (Cell signaling, 1:200), LC3AB (Cell signaling, 1:200), Olig2 (SCBT, 1:100), HB9 (DSHB, 1:200), and CASPAR-1 (A gift from Dr. Bahat MA.). Appropriate secondary antibodies were used from 488, 568 and 647 conjugated Alexa Fluor (Life Technologies). Nuclei were counter stained with DAPI (R&D). Positive cells for each marker were counted from 5 random fields of each sample with at least 3 biological repeats. For MBP staining, wells with positive fibers were counted from 10 different wells of a 24 well plates in each repeat (See also Extended Data Fig. 1).

### Fluorescence Activated Cell Sorting (FACS)

For sorting of cells, following antibodies were purchased from commercial sources: EGFR-PE (Abcam), EGFR (Abcam). Cells were harvested at different time points of the protocol from different Cell lines: day0-OCT4::GFP^+^, day8-SOX10::GFP^+^, and day 63-EGFR^+^ cells. Cell suspensions were prepared by gentle pipetting after Accutase incubation for 20 min at 37 °C. Cells were washed in ice-cold FACS buffer (2% FBS, (0.5 ug/ml) DNase I (Roche), 2 mM EDTA in PBS), then incubated with each antibody for 30 min at 4 °C and washed twice with FACS buffer. Data were acquired on a FACS MoFlo flow cytometer (DAKO) and (BD).

### RNA isolation, RNA-Seq library preparation, and sequencing

For RNA-Seq experiments, we used OCT4::GFP^+^, SOX10::GFP^+^, and EGFR^+^ sorted cells. Total RNA was isolated by TRIZOL reagent (Life Technologies), and three independent replicates were used for each population. An RNA library was prepared by JHU Deep Sequencing Core facility beginning with cDNA construction using the SMARTer pico cDNA Synthesis Kit (TaKaRa), followed by ribosomal depletion and library construction using the Illumina TruSeq Stranded Total RNA Library Prep Kit (Illumina). Sequencing was performed using the Illumina TruSeq Stranded Total RNA Library Prep Kit on an Illumina NextSeq 500 instrument using their paired-end 75 base sequencing protocol (Illumina). Sequencing yielded an average of 90.8 million reads per sample, 1.09 billion in total, with an average 83.24 mapped percentage. These reads were then aligned to the Human NCBI build GRCh38 (RefSeq June 25th,2016) using the CLCGenomicsServer 8.5.1 (with: Mismatch cost =2, Insertion cost = 3, Deletion cost = 3, Length fraction = 0.8, Similarity fraction = 0.8), comprising 3 billion bases and 53,853 identified transcripts. CLC was then used to calculate the transcripts’ FPKM (fragments per kilobase per megabase) values, which were log2 converted (FPKM = 0 ignored) and quantile normalized with Partek Genomics Suite v6.6 (Partek Inc. St. Louis USA). Known genes that demonstrated adequate expression levels, log2(FPKM) > −0.6, were used to assess differential expression between the experimental biological class groups. Differential expression analysis was carried out in Partek using the two-tailed one-way Student’s t-test algorithm, providing each gene’s differential expression (fold change) and its statistical significance (p-value). Downstream functional analyses were performed using the QIAGEN Ingenuity Pathway Analysis (IPA, QIAGEN Inc.) to determine enriched Canonical Pathways. In short, genes that showed differential expression between biological classes of >2SD, up or down, were compared to the universe of all examined genes using the Fisher’s exact test. We ranked terms according to the Fisher’s p-value, which reflect the probability of obtaining the observed overlap or greater by chance, and examined the enriched pathways for biological significance. **Data access.** The RNA-Seq data that support the findings of this study have been deposited in the NCBI database with the accession code GSE102692.

### For ZIKV infected samples

Cells were harvested 3 days post ZIKV and Mock treatment. RNA-Seq libraries were prepared from isolated total RNA, and were sequenced as previously described (PMID: 27580721). RNA-Seq libraries were generated from 1 μg of total RNA from duplicated samples per condition using the TruSeq LT RNA Library Preparation Kit v2 (Illumina) following the manufacturer’s protocol. An Agilent 2100 BioAnalyzer and DNA1000 kit (Agilent) were used to quantify amplified cDNA and to control the quality of the libraries. Illumina HiSeq3000 was used to perform 100-cycle single-read sequencing. Image processing, sequence extraction and adapter trimming were done using the standard cloud-based Illumina pipeline in BaseSpace. Single-read RNA-Seq reads were first aligned to human transcriptome annotations and genome assembly (hg19) using TopHat v2.1.1 (PMID: 23618408). The numbers of mapped reads for each condition can be found in Table SX. FPKM (<u>f</u>ragments <u>p</u>er <u>k</u>ilobase of transcript per <u>m</u>illion mapped reads) values were calculated by Cufflinks v2.2.1 (PMID: 23222703). Pairwise comparisons between infected and mock conditions were performed to detect differentially expressed (DE) genes using Cuffdiff v2.2.1 (PMID: 23222703). DE genes are defined as the ones with q-value less than 0.05. Gene ontology (GO) analyses of biological processes were performed by the Database for Annotation, Visualization and Integrated Discovery (DAVID) v6.8 (PMID: 19131956). To identify significantly enriched GO terms, a Benjamini–Hochberg procedure was used to control false discover rate (FDR) at 0.05. **Data access.** RNA-Seq data reported in this paper have been submitted to Gene Expression Omnibus (http://www.ncbi.nlm.nih.gov/geo/) with accession number GSE101919.

### RT–PCR and Real time PCR

Total RNA from each sample was isolated using TRIZOL reagent (Life Technologies), and reverse transcription was performed according to the manufacturer’s instructions of High Capacity cDNA Reverse Transcription Kit (Applied Biosystems). Real-time PCR was performed using KAPA SYBR FAST qPCR master mix (Kapa Biosystems) on an Eppendorf Realex Master cycler. Data analysis were performed by normalizing each sample CT level to its GAPDH. For RT-PCR, cDNA fragments were amplified by PCR master mix (Life Technologies) and analyzed by 1.3% agarose gel electrophoresis (for primers information see Extended Data TABEL 2).

### Western blot analysis and antibodies

Western blotting was carried out according to standard procedures. Cells were harvested on ice in Pierce IP lysis buffer and EDTA-free protease inhibitor cocktail (Roche), sonicated, and spun down briefly to clarify the lysate. Samples were separated on 4–20% mini-PROTEAN TGX gels (Biorad) and transferred to PVDF membranes. Primary antibodies used were MBP (Abcam, 1:1,000), MAG (Abcam, 1:5,000), PMP22 (BioLegend, 1:5,000), Cleaved caspase3 and LC3AB I/II (Cell Signaling, 1:1,000), CASPAR-1 (Sigma, 1:500) and GAPDH (Cell Signaling, 1:5,000) followed by horseradish-peroxidase-conjugated secondary antibodies (Cell Signaling). Densitometry measurements were carried out by ImageJ.

### *In vitro* co-culture assay

Co-culture SC and sensory neurons were 100,000 N (W6) + 200,000 SC (EGFR^+^: passage 1-3) per 22mm in M2 media for up to 3 months.

### Brazilian Zika virus preparation, titration, cell infection and plaque assay

The Brazilian strain (BR) of Zika virus was originally obtained from Fortaleza-Northeast region of Brazil strain, and corresponds to GenBank KX811222.1. To determine the viral titer, a monolayers of Vero cells (ATCC, CCL-81^TM^) were plated in 12 well plate 24 hours prior to infection. The following day, the plate was checked for confluence and the cells were infected with serial dilution of virus in OptiPro serum free media (Thermofisher) with 1% L-Glutamin (Life Technologies) rocking every 30 minutes for 5 hours at 37°C and 5% CO_2_. Post infection, cells were washed once with media and overlaid with 1/10 of 4% agarose (Gibco) in DMEM growth media (Life Technologies) containing 10% FBS (Gibco), 1% L-Glutamin (Life Technologies), 1% HEPES and 1% Pen-Strep solution (Thermofisher) for 5 days. After 5 days, 100ul of 10 mg/ml MTT (Molecular Probes) in PBS was added to the agarose monolayer and rotated to spread evenly over the surface. Plates were incubated for 3 hours at 37°C and plaques were visualized, counted, and the viral titer plaque formation unit (PFU/mL) was determined.

### Virus Infection

Zika virus infection of PNS neurons and Schwann cells were carried out at 0.4 MOI using low volume (400ul for 12 well plate) of Neurobasal media (Life Technologies) for 5-6 hours at 37°C and 5% CO_2,_ rocking plate every 30 minutes. Post infection, media was removed and washed once with Neurobasal media. Infection was carried out at 37°C and 5% CO_2_ for 3-7 days in M2 media.

### Transmission Electron Microscopy (TEM)

Samples were fixed in 2.5% glutaraldehyde, 3mM MgCl2 in 0.1 M sodium cacodylate buffer, pH 7.2 for one hour at room temperature. After buffer rinse, samples were postfixed in 1% osmium tetroxide, 0.8% potassium ferrocyanide in 0.1 M sodium cacodylate for at least one hour (no more than two) on ice in the dark. Following a DH2O rinse, samples were en bloc stained with 2% aqueous uranyl acetate (0.22 µm filtered, 1 hr, dark), dehydrated in a graded series of ethanol and embedded in Eponate 12 (Ted Pella) resin. Samples were polymerized at 37°C for 2-3 days followed by 60C overnight.

For ZIKV-infected samples were fixed with 2.5% paraformaldehyde and 1.5% glutaraldehyde, rinsed with water, after uranyl acetate was replaced with 100 mM maleate buffer, and the uranyl acetate (UA, 2%) was in 100 mM maleate. After UA, the samples were dehydrated and embedded similarly.

Thin sections, 60 to 90 nm, were cut with a diamond knife on the Reichert-Jung Ultracut E ultramicrotome and picked up with 2×1 mm formvar copper slot grids. Grids were stained with 2% uranyl acetate in 50% methanol followed by lead citrate and observed with a Philips CM120 TEM at 80 kV. Images were captured with an AMT CCD XR80 (8-megapixel camera - side mount AMT XR80 – high-resolution high-speed camera).

For quantitative analysis, myelinated fibers were counted per EM field (each 6,000 um^2^) from 15 different fields of each sample. Axon diameter and myelin thickness calculated by G-ratio calculator 1.0; Image J plug-in. A number of AVs, DMS and IMS were counted per EM field from random fields (to ensure randomization of measurements), from 3 different biological replicates.

### Cytokine array

Cell culture supernates (condition media-CM) of ZIKV and mock treated cells were harvested 3 dpi and stored at −80°c and cytokine production tested by using proteome profiler kit (Human Cytokine Array Panel A; R&D). Anti ZIKV drug tests experimental design is shown in S13A. We used Dasatinib (5uM) (Cell Signaling), Azithromycin (5uM) (Selleckchem), Emricasan (10uM) (Cayman), and Sofosbuvir (10uM) (Chem Cruz). All drugs were diluted in DMSO in a way that its overall concentration in medium was kept throughout the experiments below 0.1%.

### *In vivo* cell transplantation and imaging

All animal experiments have been conducted in accordance with protocols approved by the Institutional Animal Care and Use Committee (IACUC) at the JHU. Sciatic nerve microsurgery was carried out according to Gonzalez et al., 2014 protocol(Gonzalez et al., 2014), 1ul of (50,000 cells/ul) EGFP^+^EGFR^+^ cells were injected inside the sciatic nerve of each neonatal CD1 mice (P4) (Charles River). Sciatic nerves were isolated after 2 and 4 weeks post transplantation. Samples were fixed, cryoprotected in 30% sucrose (Sigma) in PBS, sectioned (20um) and processed for Immunofluorescence. Z-stack series were made using a Zeiss microscope (LSM700), and photographed using a monochrome digital camera (Scion Corp, USA) and ZEN software.

### In vivo ZIKV infection

To generate the microcephaly mouse model, we anesthetized pregnant ICR mice at embryonic day 13.5 (E13.5), exposed the uterine horns and injected 1 µL of ZIKV SZ01 (6.5 × 10^5 PFU/mL) or culture medium (RPMI medium 1640 basic + 2% FBS) into the lateral ventricles of embryos using a calibrated micropipette. For each pregnant dam, two-thirds of the embryos received ZIKV infection while the rest were injected with culture medium to provide littermate controls. After virus injection, the embryos were placed back into the abdominal cavity of dams and wound was closed. Tissue was collected and analyzed at P5. All experimental procedures were performed in accordance with protocols approved by the Institutional Animal Care and Use Committee at Beijing Institute of Microbiology and Epidemiology and conducted in a biological safety protection laboratory.

### Statistical analysis

Results presented as histograms are reported as mean ± s.e.m. (the mean of three in dependent experiment). *P* values for data comparison were calculated by Student’s *t*-test (GraphPad PRISM version 6.0; GraphPad Software Inc.) and were indicated by asterisks in figures unless otherwise noted.

### Data availability

All data supporting the findings of this study are available from the corresponding author upon reasonable request (including those within the paper and the supplementary).

## Results

### Modeling human peripheral myelination *in vitro*

In order to achieve *in vitro* myelination in differentiated hPSC culture, we modified a PNS neuronal differentiation protocol (Chambers et al., 2012; Lee et al., 2007) using different combinations of small molecules as a small cherry-picked compound library screening, growth factors and extracellular matrix. The culture period was extended up to 12 weeks using two different media conditions (M1/M2) in order to induce neuronal/glial differentiation along each other and myelination (Figures 1a, Sla-S1d, and Methods section). Under this condition, more than 60% of the cells were positive for BRN3a and Peripherin (peripheral neuronal markers). Myelin formation was confirmed by immunostaining with MBP, Neurofilament and CASPR-1 antibodies (Figures 1b and 1c). Transmission electron microscopy (TEM) analysis revealed the typical appearance of myelinated axons and the dense structure of compact myelin (Figures 1d-1f). Moreover, we found that in the presence of growth factor, Heregulin-1 type III (HRG), the frequency of MBP positive fibers and number of compact myelin significantly increased (Figures S1d and S1e). After 12 weeks in culture, myelinated fibers were detected at 2.31±0.83/EM field, but not in unconditioned cultures (Figures 1e). We also measured the degree of myelination derived from both human embryonic stem cells (ESCs) and induced pluripotent stem cells (iPSCs). The area-based G-factor was distributed in a 0.8 - 0.01 range (Figure 1f). These results demonstrate that, under this condition, PSCs undergo differentiation toward generating peripheral myelinated fibers *in vitro* recapitulating human PNS myelination.

To verify the existence of competent SCp in our *in vitro* myelination model, we employed FACS to isolate different cell population such as ERBB3, MAG and EGFR positive cells after 8 weeks of culture. The morphology, marker expression, differentiation and proliferation abilities of EGFR^+^ population (Lobsiger et al., 2001) are very similar to SCs (Figures 2a-2d). Moreover, 20-40% of cell population was EGFR^+^ and there were no significant differences between ESC- and iPSC-derived EGFR^+^ cells (Figure 2b). The EGFR^+^ cells could be expanded and maintained for several passages and cryopreserved. We performed RNA-Sequencing and compared the transcriptome of the EGFR^+^ cells with undifferentiated OCT4::EGFP^+^ hESCs and differentiated SOX10::EGFP^+^ neural crest (NC) (Figure 2 and S2). The hierarchical clustering of gene expression indicated that EGFR^+^ cell population is unique and completely different from undifferentiated NC cells (Figures 2e, 2f, and S2b-S2c). Compared with undifferentiated state, 4999 genes out of 53853 transcripts were identified, with 116 significantly upregulated and 105 significantly downregulated (> 2 SD, *P*_val_ < 0.05). When the EGFR^+^ population was compared to SOX10^+^ NC cells, 5426 genes out of 53853 were identified, with 131 significantly upregulated and 104 significantly downregulated (Figure S2a). These transcriptome-wide gene expression analyses showed higher expression levels of SC-related genes (Figure 2f) compared to that of PSC-related or NC-related genes (Figures S2b and S2c). These results were confirmed by immunostaining (Figure 2d) and RT-PCR (Figure S2d). For example, *POSTN*, a gene involved in SC development was highly expressed in EGFR^+^ cells, indicating their identity as a SC population. Moreover, our transcriptomic analysis of these cells revealed induction of several metabolic pathways necessary for myelination such as lipid and protein biosynthesis. Ingenuity Pathway Analysis (IPA) of differentially expressed genes indicated that ILK, WNT/β-catenin, Tec kinase, and CXCR4 were significantly activated in EGFR^+^ cells (Figures S2e-h).

**Figure 2.**
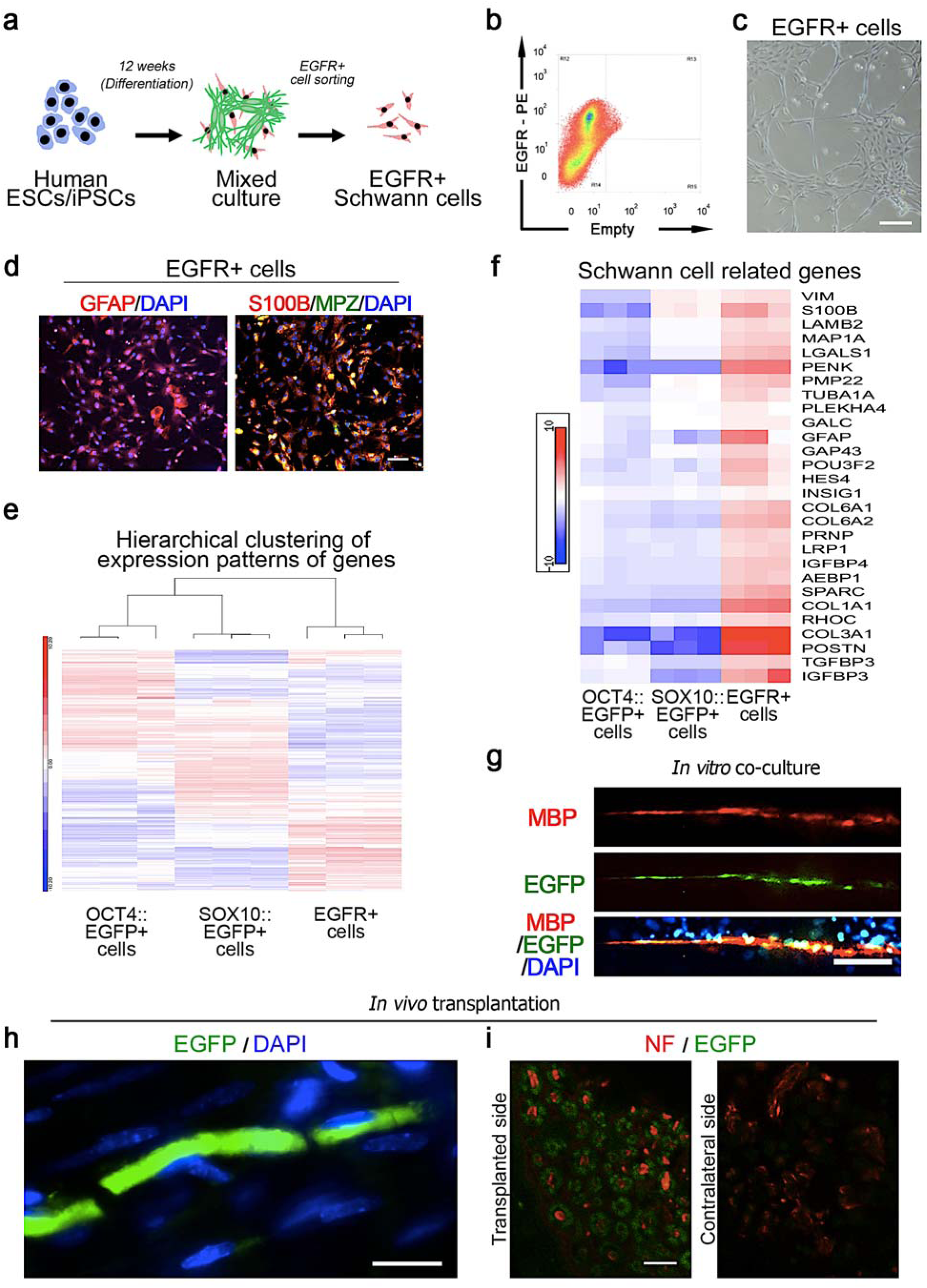
Characterization and verification of the PNS myelination model. a. Schematic summary of generation and isolation of EGFR^+^ Schwann cells from human ES/iPS cells. b. Flow cytometric representative plots of EGFR^+^ cells after 9 weeks of PNS differentiation protocol. c. Phase contrast image representative EGFR^+^ cells morphology in culture (Scale bar, 500 μm) d. Representative immunofluorescence images of EGFR^+^ cells for S100B, GFAP, and MPZ (Scale bar, 50 μm). e. Hierarchical clustering of whole-transcriptome expression profiles demonstrate wave of gene expression changes during differentiation from OCT4::EGFP (undifferentiated hPSCs), SOX10::EGFP (hPSC-NC) to EGFR positive cells, (*n =* 3 biological replicates per groups) f. Heat map of the expression levels of selected Schwann cell related genes, (*n =* 3 biological replicates per groups). g. Representative images for MBP and EGFP colocalization in the EGFR^+^EGFP^+^ cells and PNS neurons co-culture (Scale bar, 50 μm). h. Representative confocal microscope images for NF, S100 and EGFP in P4 mice pup sciatic nerve sections of EGFR^+^ EGFP^+^ cells transplanted and contralateral control sides (Scale bar, 50 μm).

Next, we sought to determine whether these cells are capable of forming myelin *in vitro* and *in vivo*. We used an EGFP expressing PSC line to demonstrate that the sorted EGFR^+^ EGFP^+^ cells could myelinate human neurons *in vitro* (Figure 2g) and mouse neurons upon transplantation inside the sciatic nerve of newborn pups (p4-around the onset of myelination) (Figures 2h). Therefore, we can conclude that the isolated EGFR^+^ cells exhibit transcriptionally distinct features and capabilities to form *de novo* myelination both *in vitro* and *in vivo*.

### Modeling human immune-mediated neuropathy *in vitro*

We utilized our *in vitro* human myelination system to model an immune-mediated neuropathy with a Brazilian ZIKV strain (ZIKV^BR^) (Figure 3a). At a multiplicity of infection (MOI) of 0.4, ZIKV could efficiently infect both PNS neurons and SCs three days post infection (3 dpi) based on immunostaining, q-PCR, and viral plaque assay (Figures 3b and S3a). These data were corroborated by intra-cranial infection studies with mice (Figure S3b). At 7 dpi, obvious characteristics of disruption of myelin sheath integrity, shrinkage, and decompaction (Figures 3c and 3d) and empty spaces with multiple intracellular oval-shaped myelin segments and debris (myelin discontinuities, myelinophagy and myelin loss) were observed by TEM analysis. In addition, broad ER-Golgi remodeling, mitochondrial damage, and increased numbers of phagosomes and lysosomes were detected (Figure S3f-S3i). The increased levels of cleaved Caspase-3 (cl-Caspase3) and LC3AB indicate the induction of cell death and autophagy by ZIKV in our myelination model (Figures 3b, and S2c-S2e). Taken together, these results demonstrate that both neurons and glia cells of PNS are targeted by ZIKV, and ZIKV infection caused cell death and destruction of myelination.

**Figure 3.**
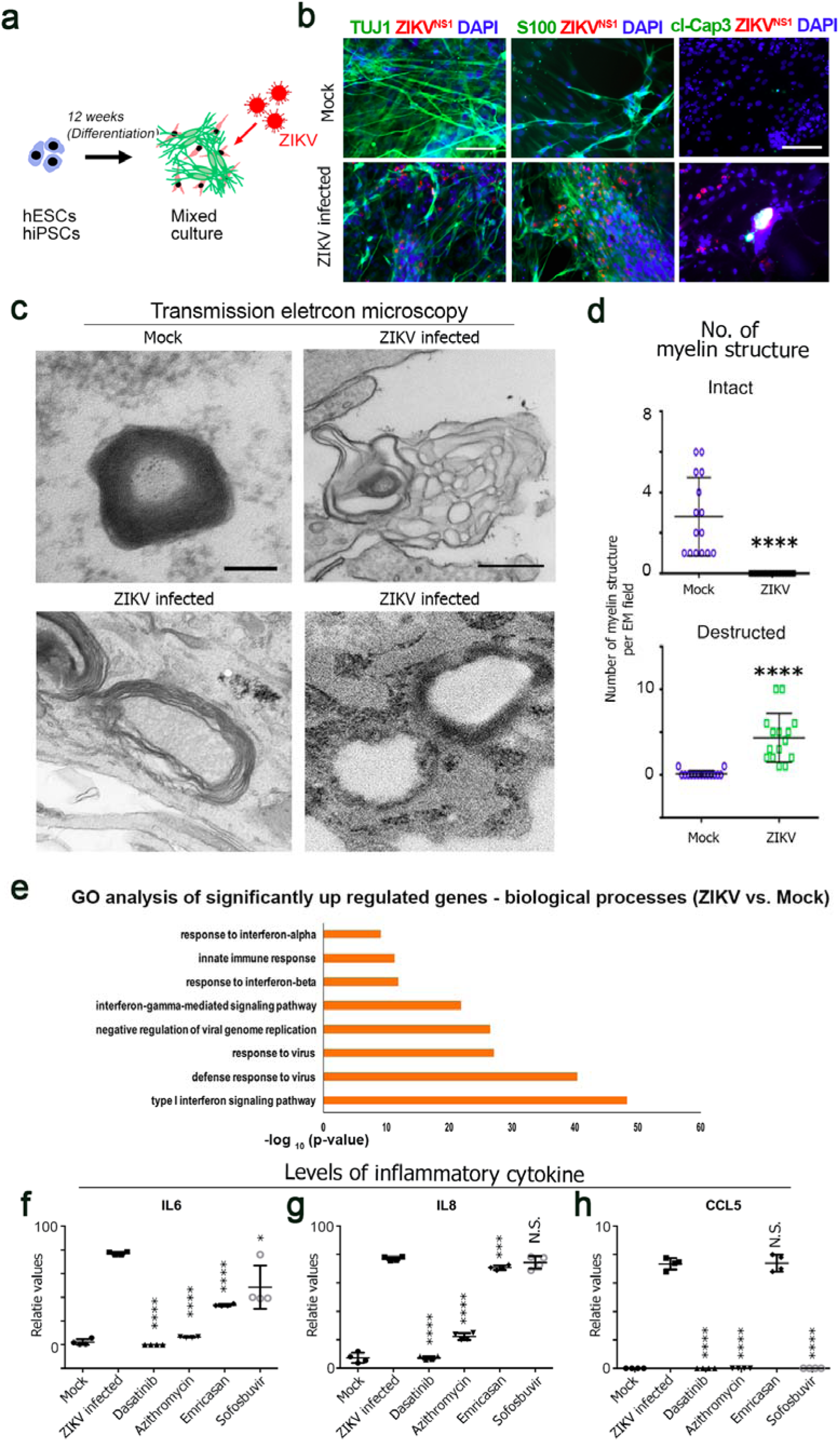
ZIKV infection induced cytokine production, demyelination and cell death. a. Schematic summary of ZIKV infection. b. Representative immunofluorescence images for ZIKV-NS1, TUJ1 or S100 and cl-Caspase3 in Mock and ZIKV^BR^ treated cultures (MOI; 0.4), 3 dpi. (Scale bar, 50 μm). c. Transmission electron microscopy (TEM) of Mock and ZIKV-infected cultures 7 dpi (Scale bar, 100 μm). illustrating the ZIKV-induced changes in myelin sheath. d. The number of intact and disrupted myelin in mock and ZIKV-treated cultures. (*n =* 15, data represent mean ± S.E.M., **** *P* < 0.0001, N.S. is non-significant, calculated by unpaired *t*-test). e. Gene ontology (GO) analyses of significantly upregulated genes identified by RNA-seq analysis of ZIKV^BR^- and mock-treated cultures 3dpi. Top enriched GO terms are shown. f-h. Quantification of cytokine array performed with human cytokine profiler panel A, of Mock- and ZIKV-treated cultures in the presence of Dasatinib (5 µM), Azithromycin (5 µM), Emricasan (10 µM) and Sofosbuvir (10 µM). (*n =* 4, data represent mean ± S.E.M., **** *P* < 0.0001, *** *P* < 0.0008, * *P* < 0.02, N.S. is non-significant, calculated by unpaired *t*-test)

We next explored the molecular fingerprints of ZIKV-infected cells by RNA-seq analysis. Significantly upregulated genes were primarily enriched in Gene Ontology (GO) terms associated with type I interferon signaling pathway (INF), defense response to virus, response to interferon-alpha and gamma, immune responses, etc. (Figure 3e delete negative regulation of viral genome replication in the fig). In addition, pathway analysis of significantly differentially upregulated genes showed functional grouped networks of viral infection, antigen processing and presentation, RIG-I-like receptor signaling (an inflammatory cytokine dependent cell death pathway), phagosome signaling, and nicotinate and nicotinamide metabolism (Figure S3j). On the other hand, gene expression significantly downregulated by ZIKV were enriched in nucleosome assembly, telomere organization, and regulators of gene expression.

To identify the paracrine factors that drive ZIKV pathogenesis, we inspected cytokine production in cell culture supernatants from three independent mock and ZIKV treated samples by cytokine array. Upon ZIKV infection, cells produced significant amounts of inflammatory cytokines including C5/C5a complement, IL6, IL8, IP-10, and CCL5 compared with controls (Figures 3F-3h, and S4b). This is in line with our finding with RNA-Seq of up-regulation of RIG-I-like receptor signaling that leads to production of these inflammatory cytokines and cell death. These results indicate that, in human PNS cells *in vitro*, ZIKV causes immune events resembling GBS-like cytokine profile (Wang, 2015). Interestingly, Dasatinib (NS4,5 blocker; inhibitor of viral induced autophagy), Azithromycin (anti-ZIKV) and Sofosbuvir (anti-ZIKV), but not Emricasan (anti-apoptosis) treatments altered the ZIKV-induced pattern of cytokine production and reduced myelin structural damages significantly (Figures 3f-3g, and S4a-S4c). These data indicate that ZIKV infection of peripheral neurons or SCs stimulate the production of cytokines, which could be partially suppressed by pharmacological intervention.

## Discussion

In this study, we developed an *in vitro* model of human peripheral myelin formation from hPSCs for the first time. We established a robust method for the isolation, expansion, co-culture and transplantation of a new population of hPSC derived myelinating SCp. To our knowledge, this is the first myelination system with SCs and peripheral neurons derived from hPSCs, amenable to disease modeling and drug tests (Clark et al., 2017; Kim et al., 2017).

We demonstrated that the competent human EGFR^+^ SCp can be readily isolated from differentiated hPSCs and form *de novo* myelination both *in vitro* and *in vivo*. The EGFR^+^ cells isolated from our *in vitro* system had unique gene expression profile including several genes and pathways that have prominent roles in different steps of SC development and myelination (Campana et al., 2003; Tawk et al., 2011) (Sonnenberg-Riethmacher et al., 2015). Importantly, these cells are expandable and cryopreserved cells reserved their myelin forming ability when co-cultured with peripheral neurons and *in vivo*.

Using our *in vitro* system, we modeled ZIKV induced neuropathy and investigated the related molecular and cellular (including ultrastructural) pathogenesis. We found that both neurons and SCs (glia cells) were direct targets of ZIKV as shown in other culture systems of both PNS and CNS (Meertens et al., 2017; Oh et al., 2017; Onorati et al., 2016; Qian et al., 2016; Souza et al., 2016). ZIKV infection destructed myelination and caused cell death in our human PNS myelination cultures. Consistent with increased numbers of GO terms and pathways involved in cytokine production by RNA-Seq analysis, cytokine array revealed higher levels of inflammatory cytokines including Complement C5/C5a, IL6, IL8, and IP-10, which are also involved in the RIG-I apoptotic and autophagy pathways (Bayless et al., 2016; Chattopadhyay and Sen, 2017). These results indicate that ZIKV infection causes immune response similar to GBS-like cytokine profiles (Nascimento and da Silva, 2017) in human PNS cells *in vitro*. Interestingly, anti-ZIKV drug treatment suppressed the ZIKV-induced cytokine production and reduced the myelin structural defects, while the anti-apoptotic agent Emricasan could not rescue ZIKV induced myelin defects (Boldescu et al., 2017; Liang et al., 2016),(Retallack et al., 2016) (Xu et al., 2016).

Taken together, our new system is of notable potential for future disease modeling, drug screening and cell replacement therapy for different human demyelination disorders and neuropathies, as well as for investigating their underlying pathogenic mechanisms.

## ACKNOWLEDGMENTS

We are grateful to Adam A. Behensky, Labchan Rajbhandari, Feiran Zhang, Yohan Oh, Dezun Ma, Stephen M. Eacker, Qing-Feng Wu, Hongjun Song, Guo-Li Ming, Peng Jin, Arun Venkatesan, Ahmet Hoke and Gabsang Lee for all of their wonderful support and guidance for this project. This work was supported by grants from Maryland Stem Cell Research Funding (MSCRF-TEDCO) for Postdoctoral Fellowship (F.M.), the Robertson Investigator Award from New York Stem Cell Foundation (G.L), Maryland Stem Cell Research Funding (MSCRF; G.L., H.S., G.M.), start-up funding from IGDB of CAS and China NSF (31771131 to Q.W), NIH (R37NS047344, U19MH106434 to H.S, R35NS097370, U19AI131130 to G-l.M. and R01NS093213 to G.L.). The authors (G.L., Y.O., G.M., H.S.) acknowledge the joint participation by the Adrienne Helis Malvin Medical Research Foundation. We thank H. Zhang at Flow Cytometry Core Facility (JHSPH), B. Smith and M. Delannoy at Microscope Core Facility (JHU-MicFac) and C. Talbot at Deep Sequencing Core Facility (JHU) for their kind supports. We appreciate Prof. M. A. Bhat (University of Texas Health Science Center, Texas, USA) for the anti-CASPR-1 antibody

## Author contribution

F.M., and Z.X.: conception and study design, performing experiments, data analysis and interpretation, data assembly, and interpretation of data, and writing manuscript. C.Q., providing of ZIKV and interpret of data.

## Supplemental figure legends

**Supplemental Figure 1. Development of an *in vitro* human PSCs derived PNS myelination model; protocol set up, Related to Figure 1**.

a. Schematic overview of compound screening for *in vitro* PSC-PNS myelination.

b. Quantification of the number of MBP positive neurons per well from compound screening. We made a small library consist of about 38 compounds, drug was added at final concentration as in Extended Data TABEL 1, every other day for the myelination phase of the protocol (from week 6 to 12). Negative control (unconditioned cultures) were prepared with the same time exposure and amount of DMSO (Small molecules were diluted in DMSO in a way that its overall concentration in medium was kept throughout the experiments below 0.1%) (*n =* 30 wells of 24 well plates, data collected from at least 3 different independent experiments).

c. Quantification of the percentage of wells with at least one MBP positive neurons out of 30 repeats of compound screening.

d. The number of wells with at least one posit 30).

e. The number of compact myelin per EM field in presence and absence of (+/-) HRG (*n =* 15) Data in c-d represent mean ± S.E.M., **** P < 0.0001, N.S. is non-significant, calculated by unpaired t-test. Representative TEM image of compact myelin (Scale bar, 100 μm).

**Supplemental Figure 2. Characterization and identification of EGFR^+^ cells, Related to Figure 2**.

a. Volcano plots of up and down regulated differentially expressed genes in EGFR^+^ cells compared to the control groups (OCT4::GFP^+^ and SOX10::EGFP^+^ cells) in RNA-Seq analysis (P<⍰ 0.05, *n =* 3 biological replicates per groups).

b. Heat map of selected PSC-related genes and c. NC-related genes, (*n =* 3 biological replicates per groups).

d. RT-PCR analysis of SC-related genes in EGFR^+^ cells compared to the controls.

e. Ingenuity pathway analysis (IPA) for EGFR^+^ cells compared to OCT4^+^ and F. SOX10^+^ cells.

g. Hierarchical clustering of whole-transcriptome expression profiles of unconditioned and M1/M2 treated EGFR^+^ cells after 12 weeks.

h. Ingenuity pathway analysis (IPA) for EGFR^+^ cells in M1/M2 culture condition over unconditioned.

**Supplemental Figure 3. ZIKV infection, Related to Figure 3**.

a. The level of ZIKV genomic RNA replication in Mock and ZIKV-infected cultures 7 dpi.

b. Representative images of sciatic nerve sections stained with ZIKV, GFAP and Tuj1 antibodies from ZIKV-infected animals. Scale bar, 50 μm.

c. Quantification of cl-caspase3 immunostaining, 3 dpi.

d. Western blot analysis of ZIKV-NS1, Cl-Caspase and LC3ABI/II, 3 dpi.

e. Quantification of Cl-Caspase and LC3ABI/II western blotting in Mock and ZIKV-infected cultures.

f. Representative images of ER-Golgi ZIKV-induced remodeling 7 dpi compared to Mock (Scale bar, 500 μm, higher magnification of boxed region illustrating remodeled ER -rER) and right-side image demonstrates ZIKV virions (ZVi), damage to mitochondria (Mito) and boxed region represents virus-induced vesicles (Ve) (Scale bar, 100 μm).

g. Myelinophagy by Schwann cells (Scale bar, 100 μm).

h. Representative images of ZIKV increased autophagic vesicles (AVs), and lysosome, 7dp.i. (Scale bar, 100 μm).

i. Quantification of autophagic vesicles in Mock and ZIKV-infected cultures 7dp.i. **** P < 0.0001.

j. Subset-heat map for expression levels of genes involved in RIG-I and Phagosome pathways in Mock and ZIKV-infected cultures.

**Supplemental Figure 4. ZIKV-induced defects on myelin sheath, Related to Figure 3**.

a. Schematic illustration of anti-ZIKV drug test on 12 week-cultures of hPSC-PNS *in vitro* model, condition media-CMs-were harvested 3 dpi for cytokine profiling.

b. Quantification of cytokine production from Mock, ZIKV and drug treated ZIKV-infected cultures, (*n =* 2-5 biological repeats per each condition, data represent mean ± S.E.M., **** P < 0.0001, N.S. is non-significant, calculated by unpaired t-test).

c. The number of intact or destructed myelinated fibers per EM field in presence and absence of ZIKA, and ZIKA plus drugs (Mock, ZIKV infected, and ZIKV+ Dasatinib, +Azithromycin, +Emricasan, +Sofosbuvir) (n = 15 biological repeats per each condition, data represent mean ± S.E.M., **** P < 0.0001, N.S. is non-significant, calculated by unpaired t-test).

**Supplemental TABLE 1.** List of chemical compounds and their concentration used for protocol set up.

| Small molecules | Concentration | Company |
| --- | --- | --- |
| Clobetasol | 5uM | Cayman |
| Miconazole | 5uM | Cayman |
| Clemastidine | 1uM | Cayman |
| Rapamycin | 25nM | Santa Cruz |
| Staurosporine | 1uM | Santa Cruz |
| U46619 | 100nM | Santa Cruz |
| Celastrol | 0.5uM | Cayman |
| Calphostin C | 1uM | Cayman |
| GANAT61 | 1uM | Cayman |
| GANAT58 | 1uM | Cayman |
| Anisomycin | 1uM | Cayman |
| SF1670 | 3uM | Cayman |
| PI3K activator | 1uM | Cayman |
| Biotin | 250uM | Cayman |
| PP2 | 10uM | Cayman |
| B-estradiol | 1uM | Cayman |
| Heparin | 1uM | Sigma |
| DHEA | 1uM | Sigma |
| CollageneVI | 10ug | Sigma |
| WNT1 | 50nM | Peprotec |
| WNT3A | 50nM | R&D |
| WNT7A | 60nM | Sigma |
| WIP | 50nM | R&D |
| XAV939 | 5uM | Cayman |
| PMP | 250ng | Sigma |
| SHH | 5uM | R&D |
| Cyclopamine | 10nM | Sigma |
| BMP4 | 25ng | R&D |
| TGFb1 | 10uM | R&D |
| DAPT | 10nM | Cyman |
| PD | 3uM | Sigma |
| FGF2 | 2ng | R&D |
| CHIR | 3uM | Tocris |
| ROCK inhibitor | 1uM | Cyman |
| Compound C | 1uM | Millipore |
| 8-Bro cAMP | 500uM | Sigma |
| 5AZAC | 0.5uM | Sigma |
| TSA | 5ng | Sigma |
| VPA | 500uM | Sigma |
| DMSO | >0.1% | Sigma |

**Supplemental TABEL 2.**
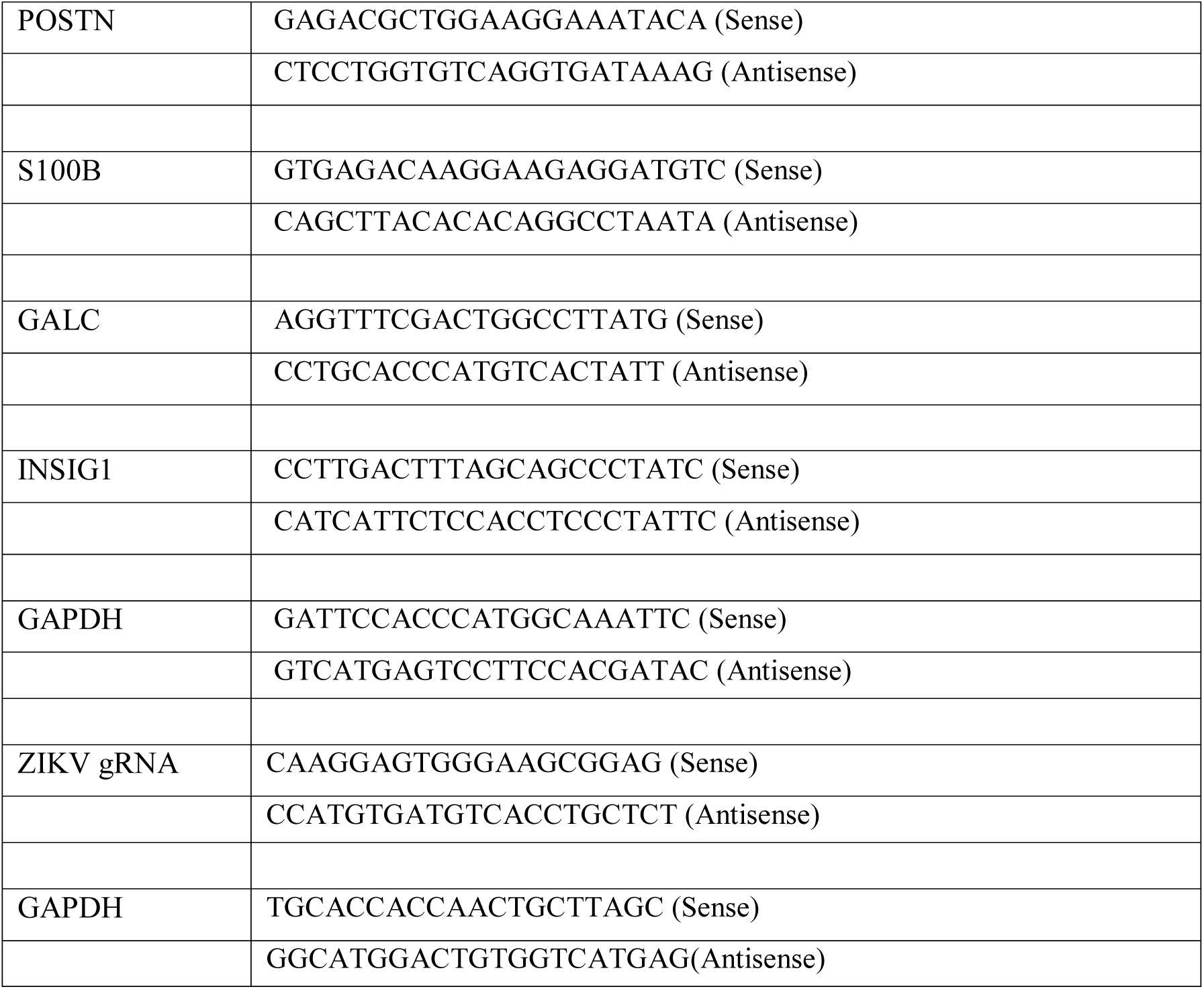
The q-PCR and RT-PCR primer sequences.

## Notes

### Competing Interest Statement

The authors have declared no competing interest.

