## Supplementary material for "Modeling Human Peripheral Myelin Fully from Pluripotent Stem Cells and Immune-mediated Neuropathy *In Vitro* using ZIKA Virus": Suppl figures

Mirakhori, et al. **Supplemental figure 1.**

Mirakhori, et al. **Supplemental figure 1.**

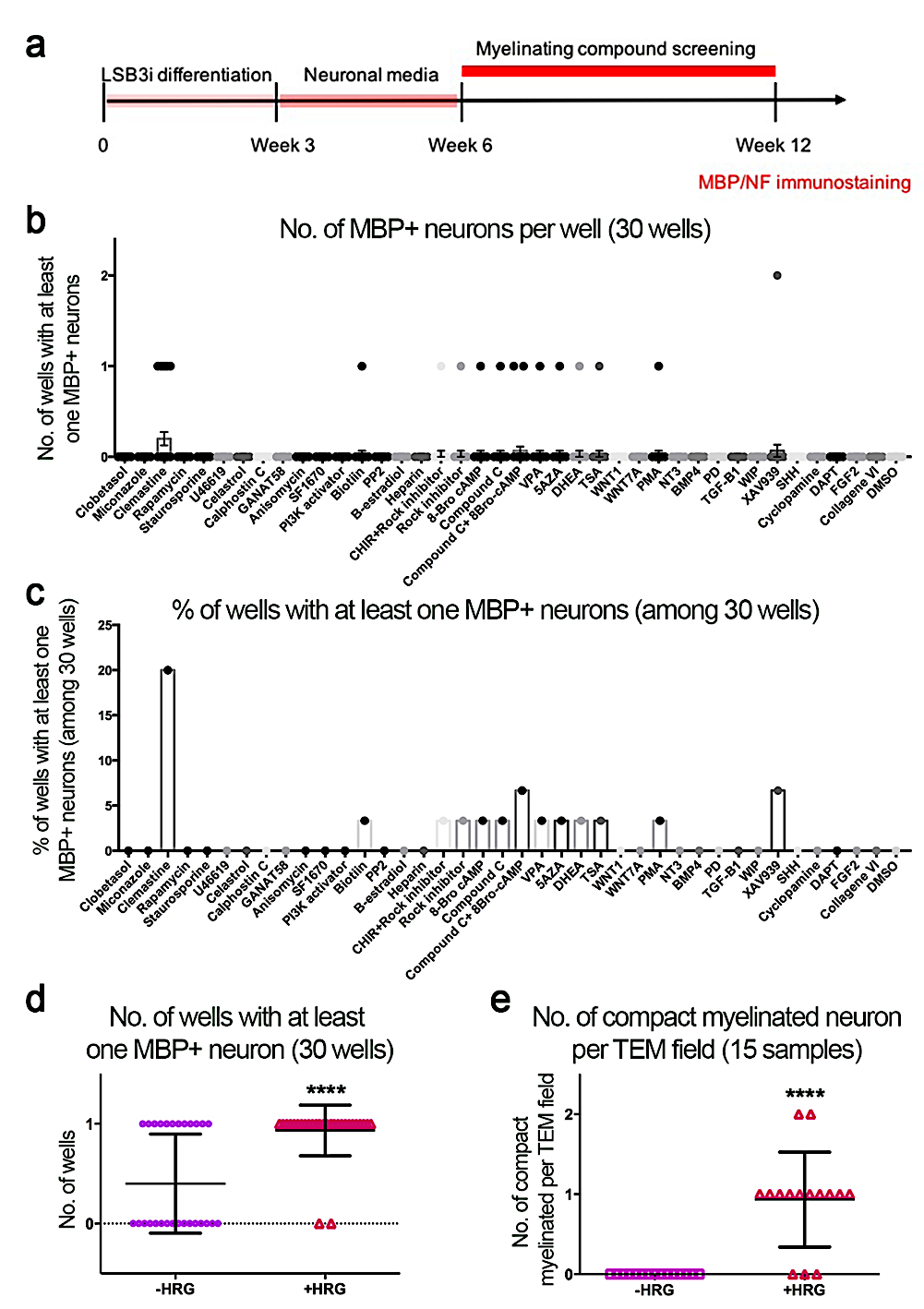

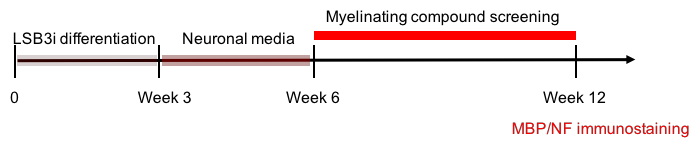

Myelination media +/- HRG

Immunostaining, TEM

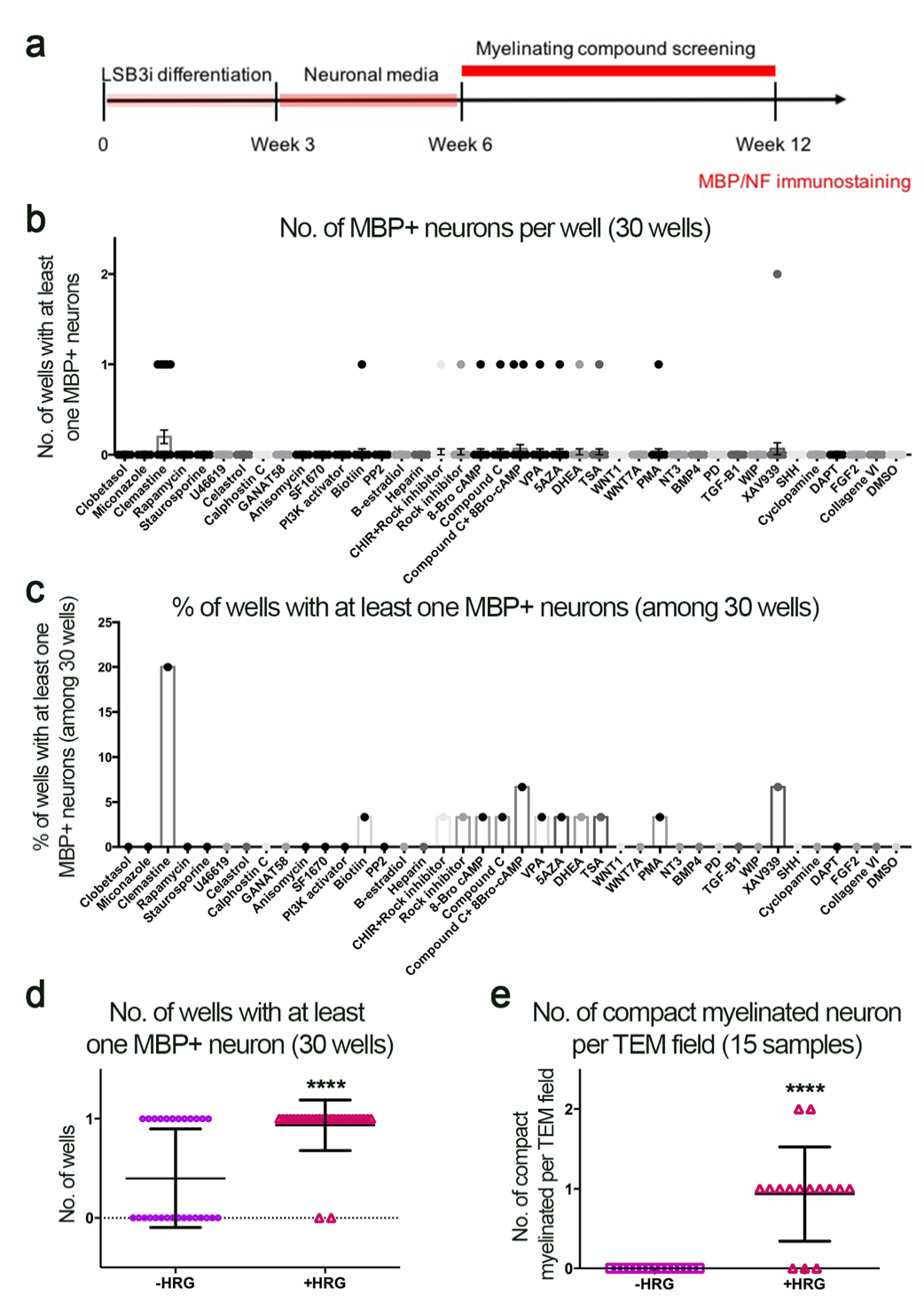

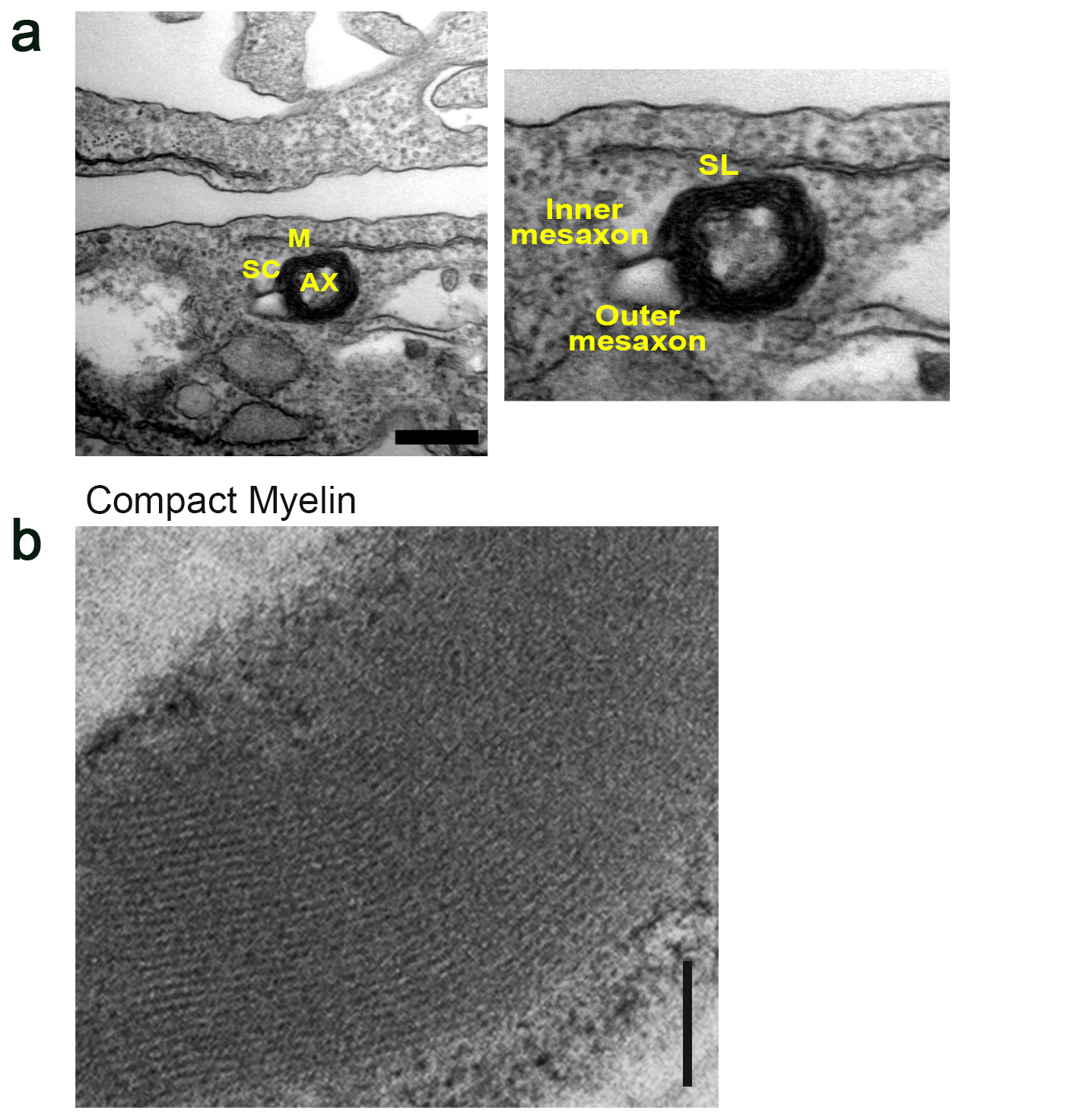

Mirakhori, et al. **Supplemental figure 2.**

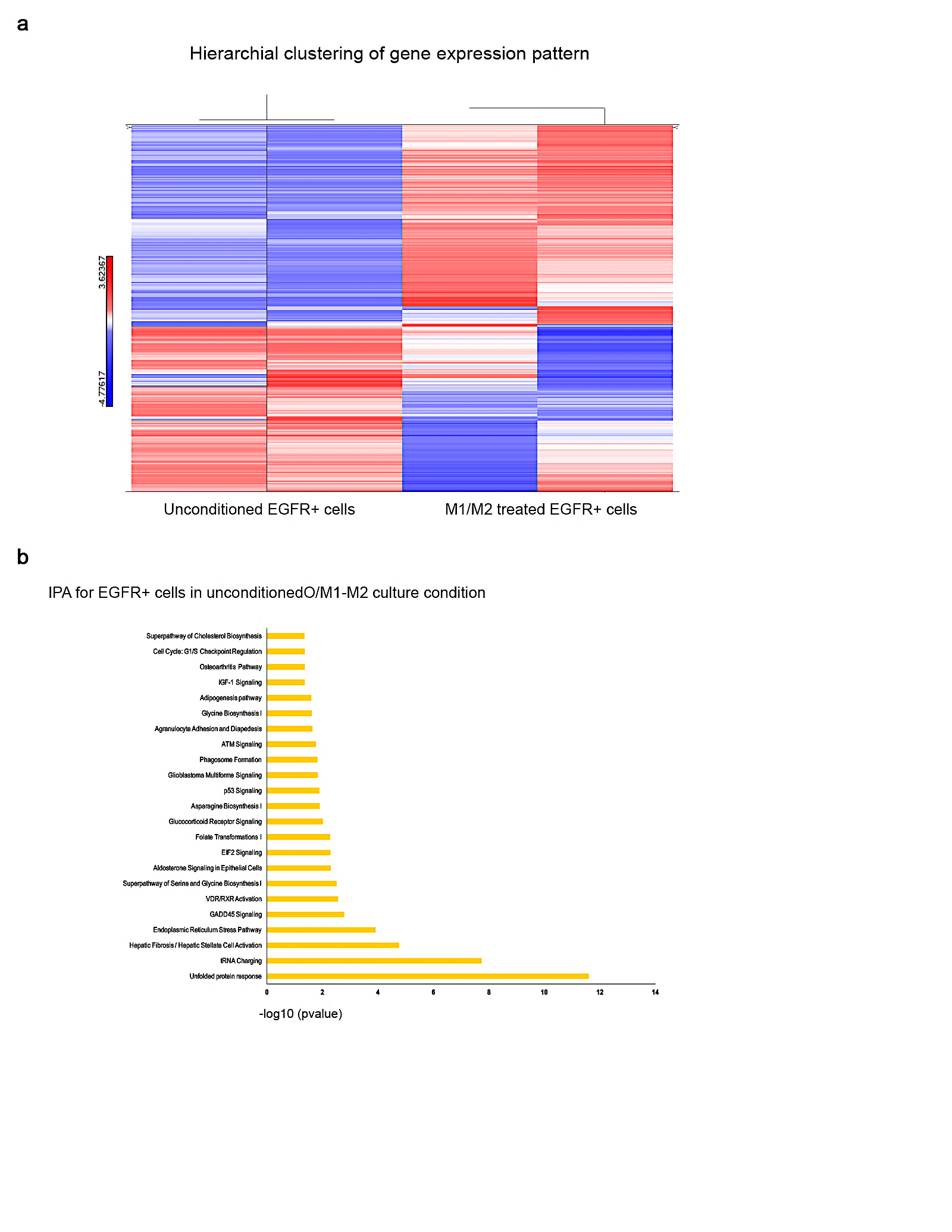

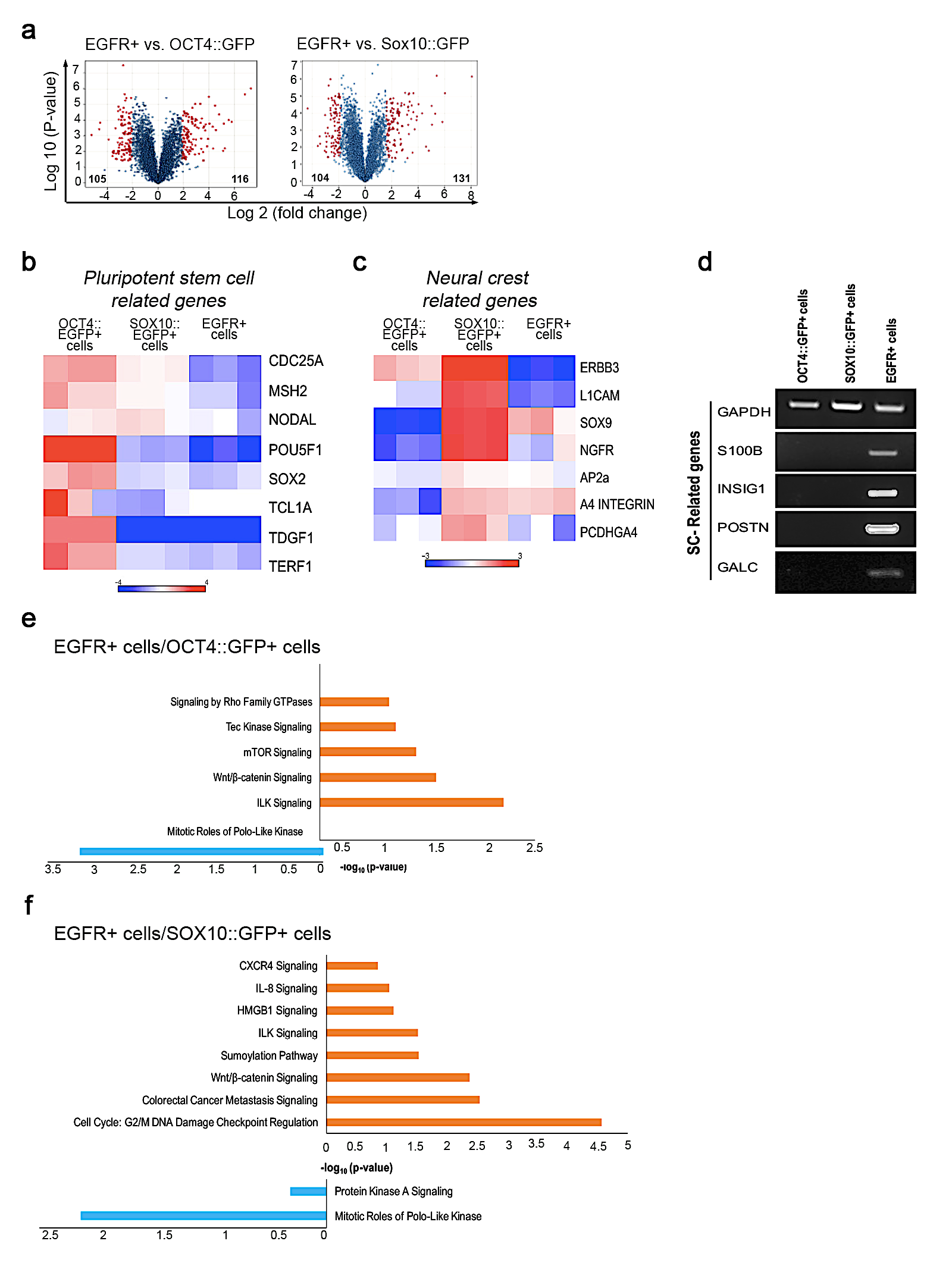

**h**

Hierarchical clustering of gene expression pattern

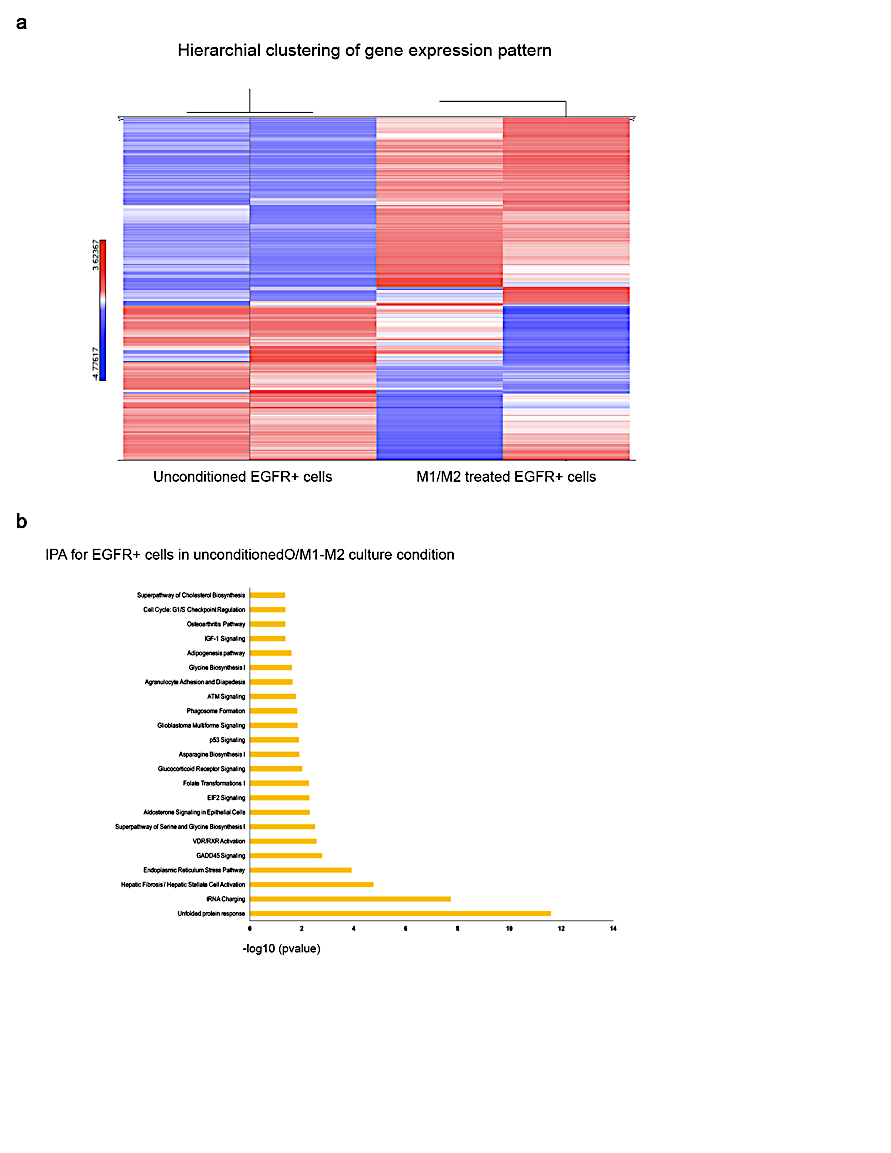

**g**

IPA for EGFR+ cells under M1/M2 condition expression pattern

Mirakhori, et al. **Supplemental figure 2.**

Mirakhori, et al. **Supplemental figure 3.**

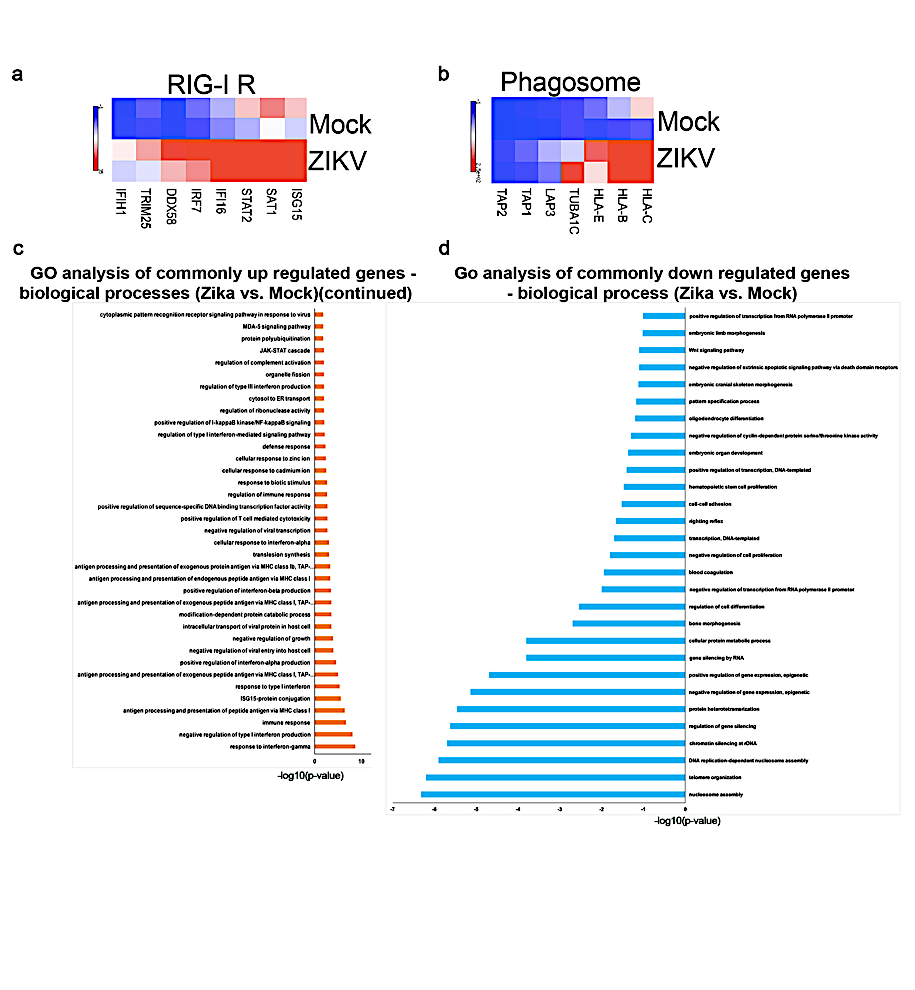

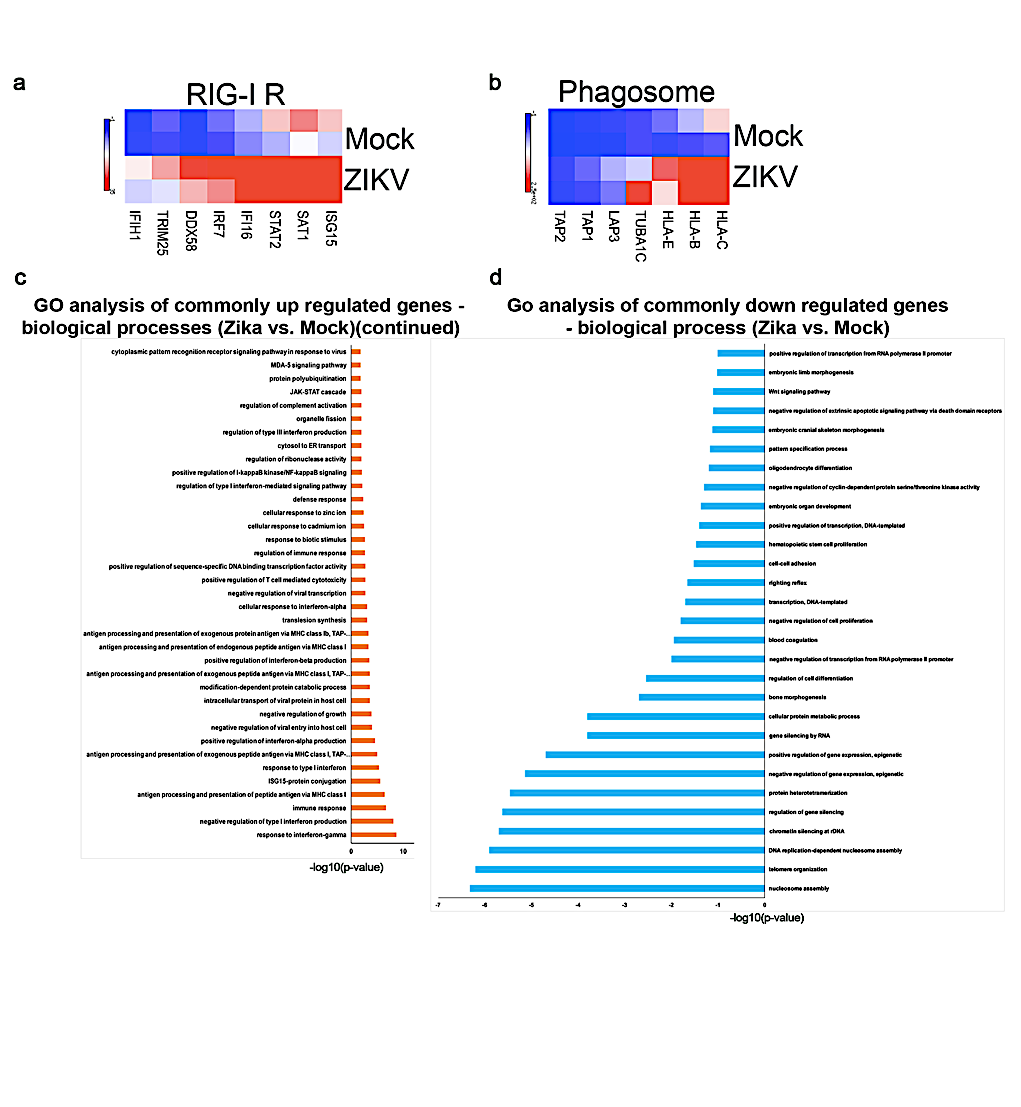

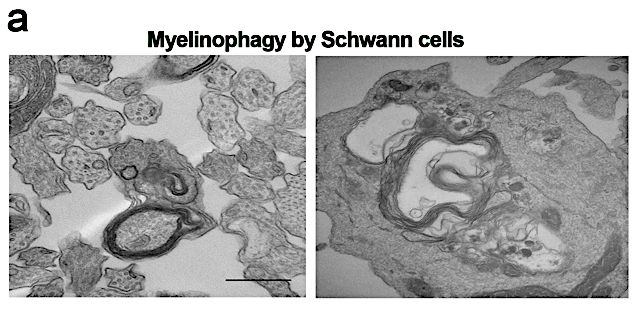

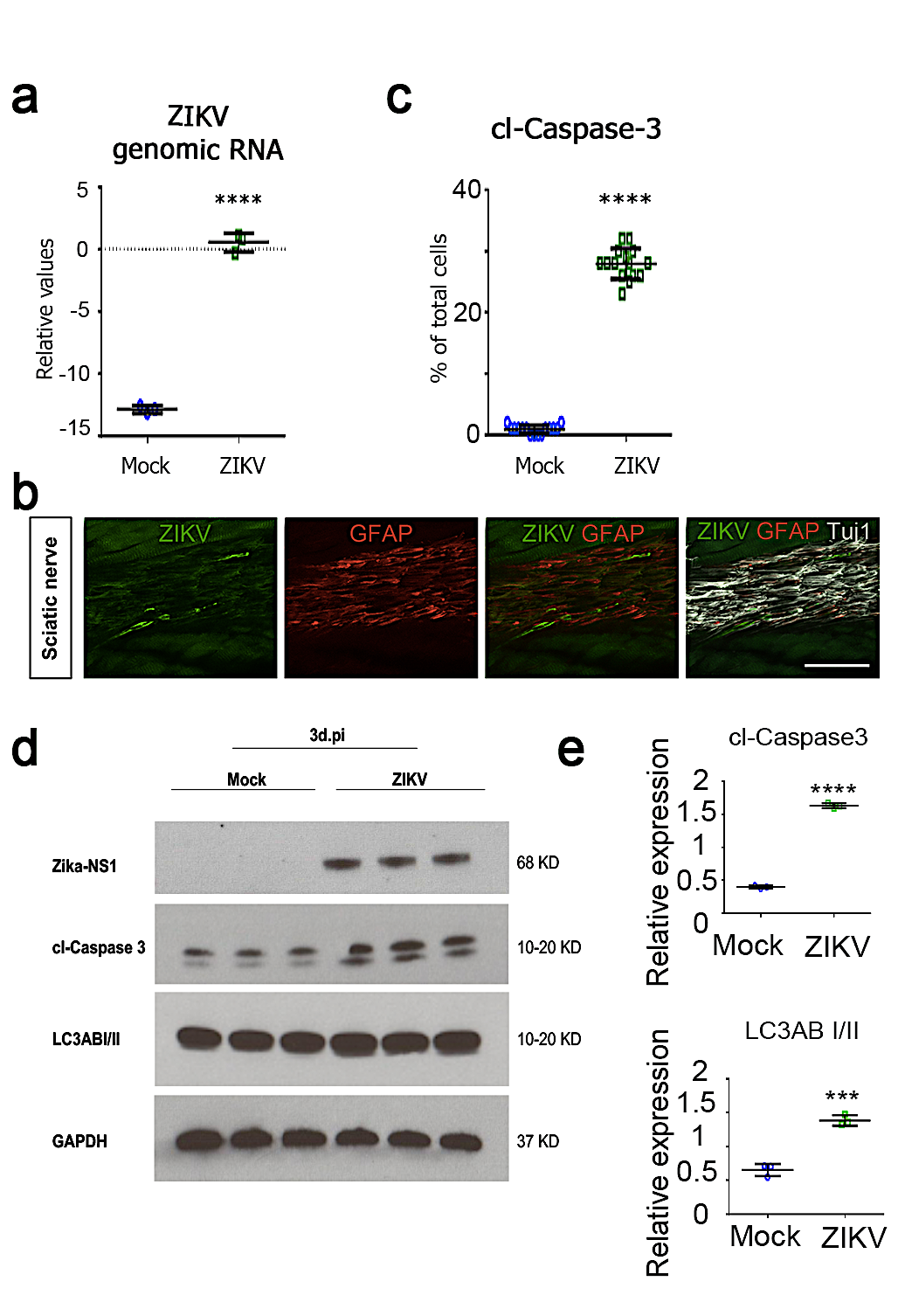

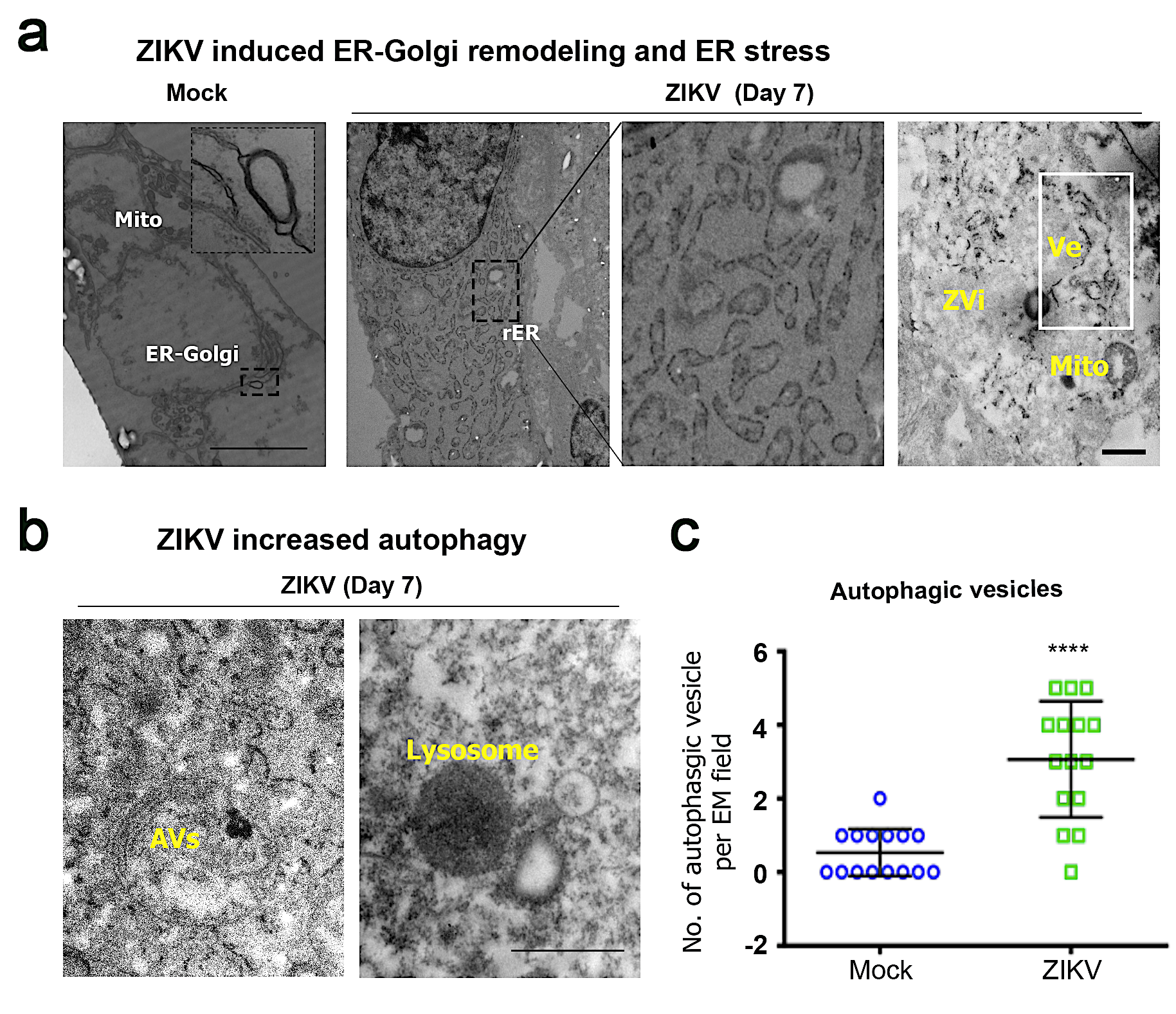

j

i

h

g

f

Mirakhori, et al. **Supplemental figure 3.**

Mirakhori, et al. **Supplemental figure 4.**

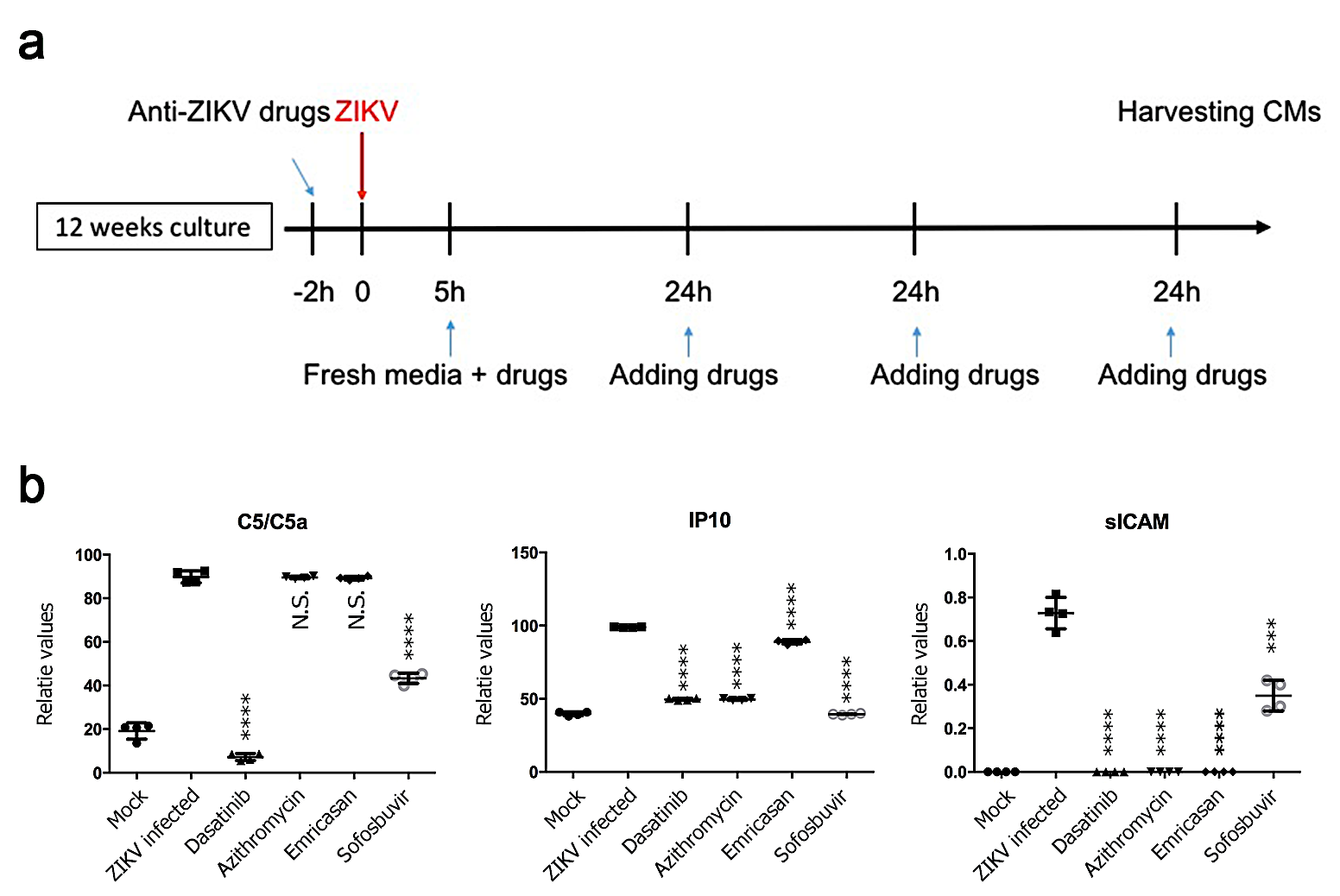

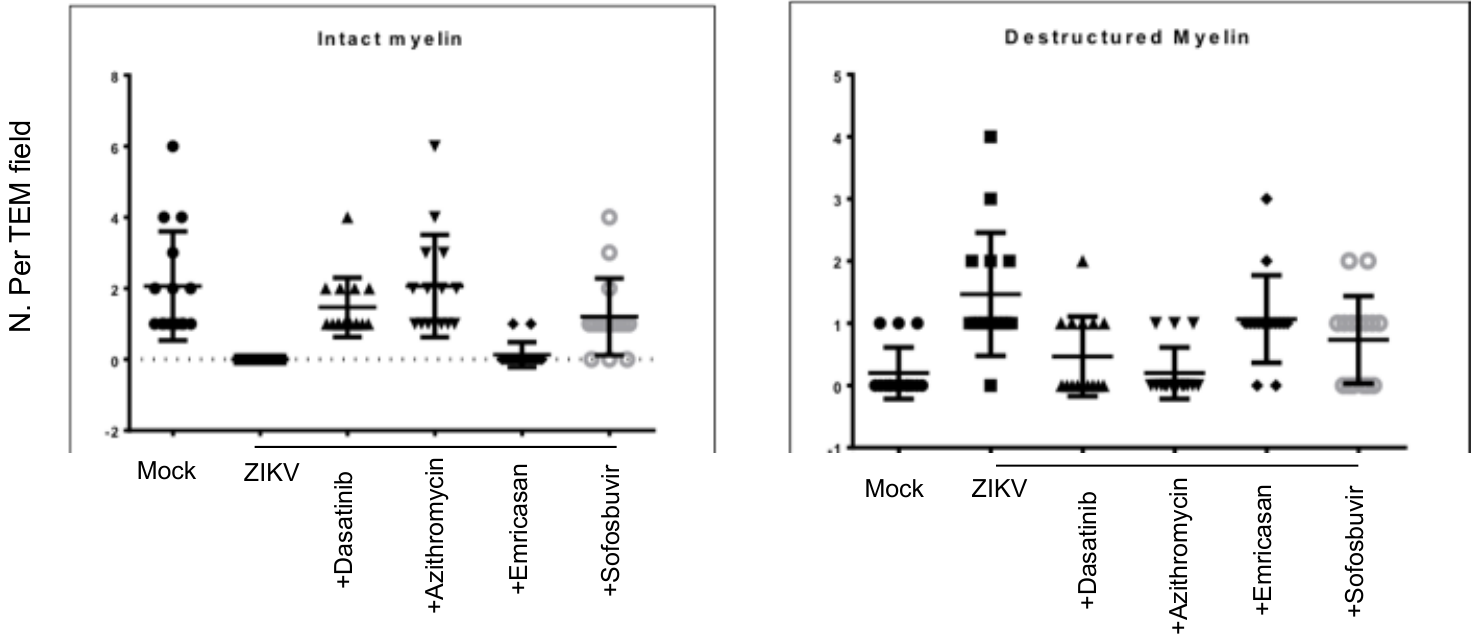
 c

**C**

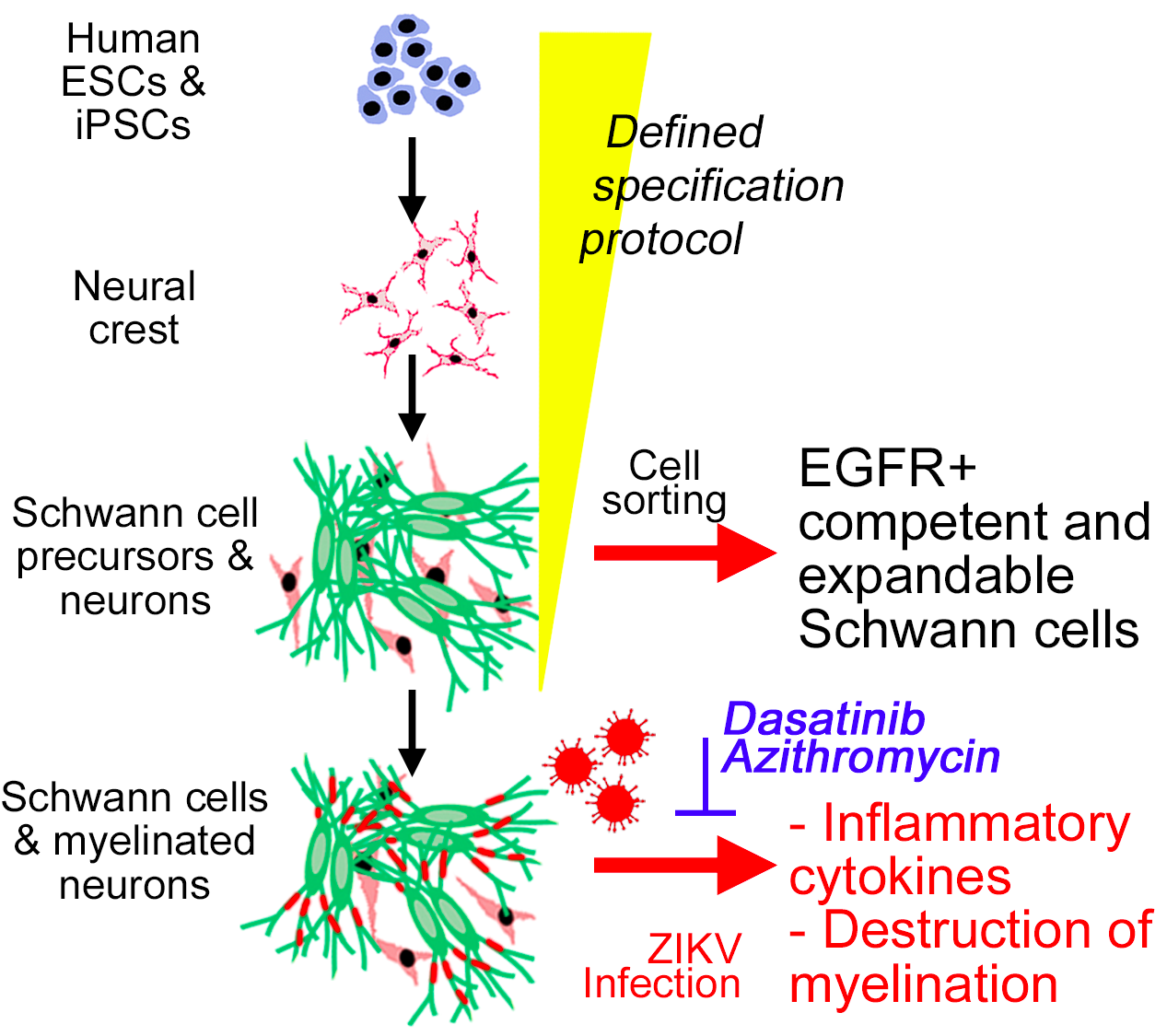

Mirakhori, et al. **Graphical abstract**
